# A microwell cell capture device reveals variable response to dobutamine in isolated cardiomyocytes

**DOI:** 10.1101/636852

**Authors:** J. A. Clark, J. D. Weiss, S. G. Campbell

## Abstract

Isolated ventricular cardiomyocytes exhibit substantial cell-to-cell variability, even when obtained from the same small volume of myocardium. In this study, we investigated the possibility that cardiomyocyte responses to *β*-adrenergic stimulus are also highly heterogeneous. In order to achieve the throughput and measurement duration desired for these experiments, we designed and validated a novel microwell system that immobilizes and uniformly orients isolated adult cardiomyocytes. In this configuration, detailed drug responses of dozens of cells can be followed in parallel for intervals exceeding one hour. At the conclusion of an experiment, specific cells can also be harvested via a precision aspirator for single-cell gene expression profiling. Using this system, we followed changes in Ca^2+^ signaling and contractility of individual cells under sustained application of either dobutamine or omecamtiv mecarbil. Both compounds increased average cardiomyocyte contractility over the course of an hour, but responses of individual cells to dobutamine were significantly more variable. Surprisingly, some dobutamine-treated cardiomyocytes augmented Ca^2+^ release without increasing contractility. Other cells responded with increased contractility in spite of unchanged Ca^2+^ release. Single-cell gene expression analysis revealed a significant correlation between expression of PRKACA and cellular sensitivity to dobutamine, demonstrating that variable drug responses among cells are not merely experimental artifacts. By enabling direct comparison of the functional behavior of an individual cell and the genes it expresses, this new system constitutes a unique tool for interrogating cardiomyocyte drug responses and discovering the genes that modulate them.

**SIGNIFICANCE:** We have created a microwell capture device that allows drug responses of dozens of isolated adult cardiomyocytes to be monitored for extended intervals. Using the device, we observed striking diversity in the Ca^2+^ handling and contractility responses of each cell to a *β*-adrenergic agonist. This included cells that responded to dobutamine by doubling the amplitude of Ca^2+^ release while decreasing contractility and vice-versa. We further show that these diverse responses can be linked to gene expression differences between cells. This work demonstrates the feasibility of linking the drug responses of individual cells with their gene expression. It opens the possibility of exploiting cell-to-cell variation to discover new genes that participate in and modulate regulatory cascades.

## INTRODUCTION

Individual isolated cardiomyocytes display an astonishing degree of functional heterogeneity. We recently reported that cells isolated from the same myocardial region vary in their intrinsic relaxation rate by more than 25% of the population mean (1). At least part of this variability can be explained by troponin I (TnI) phosphorylation, which was found to be elevated in the most rapidly-relaxing cells. Since TnI phosphorylation is a known endpoint of the cardiac *β*-adrenergic signaling pathway, our observations provide implicit evidence of cell-to-cell variation in this regulatory cascade.

Seeking direct evidence to support this hypothesis, we studied single-cell responses to dobutamine, a *β*-adrenergic agonist and potent positive inotrope often used in clinical settings (2). Omecamtiv mecarbil (OM), another positive inotrope (3), was selected as an adrenoceptor-independent control. Unlike dobutamine, which ultimately affects many intracellular targets, OM is a direct small-molecule activator of cardiac myosin with high molecular specificity. We therefore expected cardiomyocyte contraction to respond more uniformly to OM than to dobutamine.

To facilitate these studies, we designed and validated a novel device that traps cardiomyocytes into a grid of micro-patterned wells. These wells provide a uniform orientation and consistent positioning of isolated cardiomyocytes, even during fluid flow. The device enables the rapid and robust measurement of dozens of cells in the same experiment, with the ability to track cells during one hour, or more, of drug exposure.

By coupling this microwell array with a precision aspirator (1), it was also possible to collect specific cells after an experiment and perform single-cell gene expression assays. To our knowledge, the experiments reported here constitute the first successful attempts to relate a single-cardiomyocyte drug response to gene expression in the same cell. The resulting data expose surprising cell-to-cell variation in dobutamine responses and provide evidence that adaptation to sustained *β*-adrenergic stimulation is also cell-specific.

## MATERIALS AND METHODS

### Ethical approval

All animal procedures were approved by the Yale University Institutional Animal Care and Use Committee (Approval #2015-11528), compliant with the regulations of the Animal Welfare Act, Public Health Service, and the United States Department of Agriculture. Studies were performed using 9 female Sprague Dawley rats, comprising retired breeders 4-6 months of age purchased from Charles River Laboratories (Wilmington, MA). Animals were housed under a standard 12:12 h light/dark cycle and fed ad libitum in accordance with an approved Yale University protocol. Animals were exposed to 15 min of 500 mL min-1 isoflurane inhalation, then were subjected to a terminal bilateral thoracotomy for removal of the heart. All experiments were conducted in accordance with the relevant guidelines established standards.

### Cardiomyocyte isolation

Left ventricular adult rat cardiomyocytes were isolated via Langendorff perfusion of the heart with a collagenase-containing solution. Adult female rats were anaesthetized with isoflurane for 15 min, injected with 0.5 mL of 1000 U mL^−1^ heparin, and subjected to bilateral thoracotomy for removal of the heart. Excised hearts were cannulated via the aorta and mounted onto a Langendorff perfusion apparatus within 3 min of excising the heart. Hearts were immediately perfused for 15 min with 37°C calcium-free perfusion buffer containing (in mM): 118 NaCl, 4.8 KCl, 1.25 KH_2_PO_4_, 1.25 MgSO_4_, 10 2,3-butanedione monoxime, 25 Hepes, and 11 glucose, supplemented with 10 μM (±)-Propranolol (hydrochloride) (Cayman Chemical, Ann Arbor, MI) (pH 7.3). A digestion buffer was next perfused through the heart for 20 min. The digestion buffer consisted of perfusion buffer supplemented with (in mM): 2.5 carnitine, 5 taurine, 2.5 glutamic acid, 0.025 CaCl_2_, 120 U mL^−1^ collagenase type II (Worthington Biochemical Corp., Lakewood, NJ), and 0.96 U mL^−1^ protease (Sigma-Aldrich, St. Louis, MO) (pH 7.3). The heart was then cut down from the apparatus and the left ventricular free wall was removed. The sub-epicardial portion of this left ventricular excision was cut free and the innermost third of that sub-epicardial layer was isolated to avoid edge effects from the base and apex. The resulting strip was segmented into three equal regions of ~1 x 1 x 1 mm (length x width x thickness) in size per region. Each ~1 mm^3^ region then went on to be digested through mechanical agitation on a shaking incubator set at 115 r.p.m. for 10 min at 37°C and then gently triturated to liberate individual cells from the tissue. Undigested tissue chunks were then transferred to fresh digestion buffer in a separate microcentrifuge tube, and the process was repeated as many as three additional times, or until all tissue had been digested. Cells were removed from digestion buffer by centrifugation and resuspended in a sequence of perfusion buffers supplemented with fetal bovine serum and with gradually increasing calcium concentrations (0.05-1.1 mM). Functional experiments were performed after allowing cells to rest for at least one hour.

### Microwell device and transcriptomics overview

In order to measure sarcomere length changes and Ca^2+^ transient records from Fura-2-loaded cardiomyocytes consistently over multiple time points, we designed a custom microwell polydimethylsiloxane (PDMS) substrate within a 3D printed cell bath that kept the cells uniformly oriented (Fig. 1A). An array of rectangular posts allowed cells to be exposed to superfused physiological buffer and electrical stimulus from platinum pacing electrodes while preventing them from drifting away from the previous measurement window (Fig. 1B and 1C). After taking functional records, some of the cardiomyocytes were individually aspirated and deposited via an automated single-cell aspirator into microcentrifuge wells filled with lysis buffer for transcriptomic analysis.

**Figure 1:**
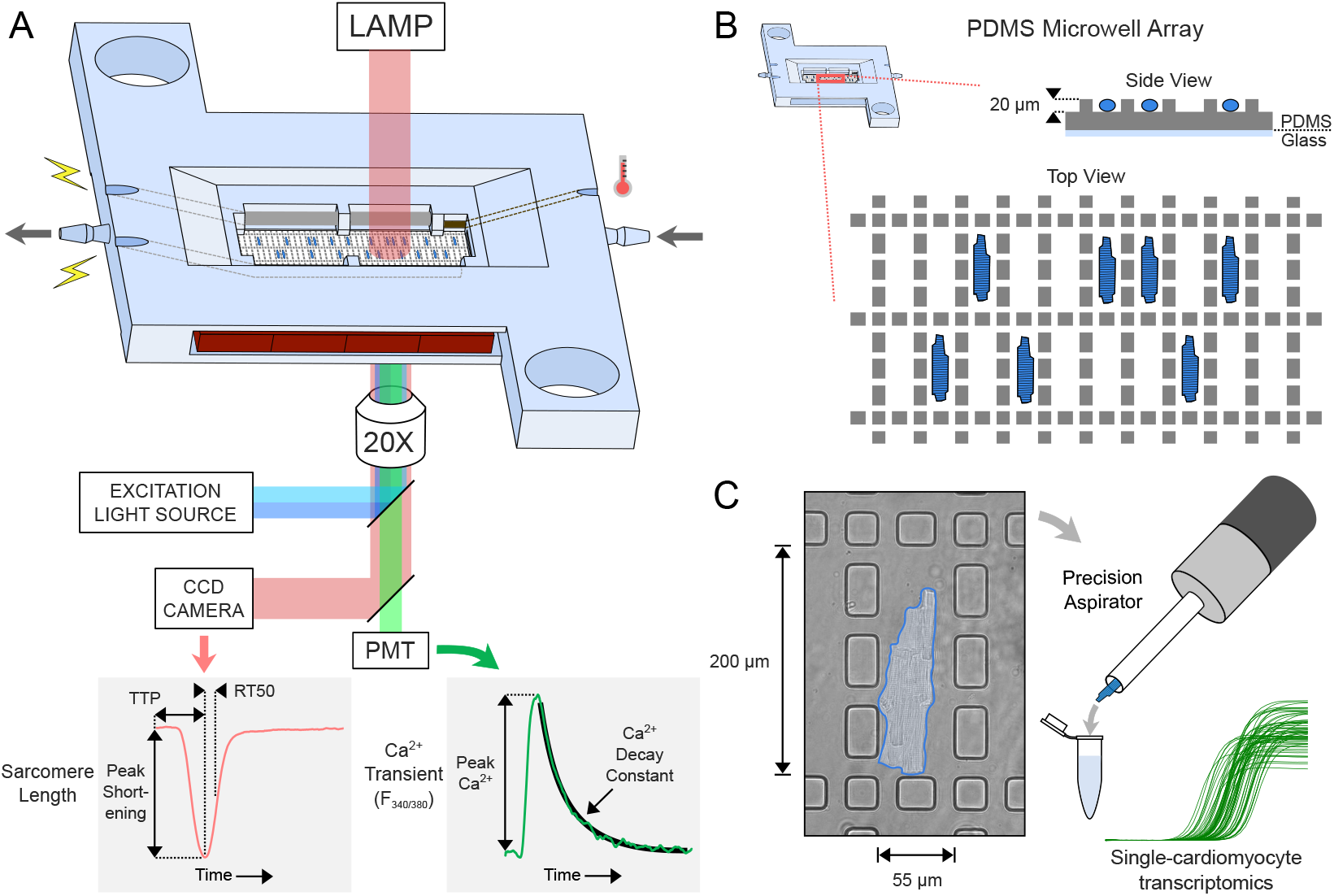
Overview of experimental apparatus and PDMS microwell device for cell capture and transcriptomics. *(A)* Schematic of device with optical data collection system below. Arrows indicate direction of fluid flow. Device contains platinum pacing electrodes, heating elements, and a temperature sensor. *(B)* Enlarged diagram of PDMS microwell pattern. Each filled rectangle is a pillar between which cells are captured. *(C)* Bright-field micrograph (left) of a captured cardiomyocyte, outlined in blue, under 30x magnification. Cells can be collected with an automated single-cell aspirator (right) for single-cardiomyocyte transcriptomic analysis.

### PDMS microwell manufacturing

SYLGARD 184 Silicone Elastomer Base and Curing Agent (Dow Chemical, Midland, MI) were mixed thoroughly in a 8:1 mass ratio and vacuum desiccated for 20 min until all air bubbles were removed. The resulting solution was spin coated (Laurell Technologies, North Wales, PA) on a SU8 silicon master wafer (soft lithography negative photoresist) containing the microwell pattern at 400 RPM for 60 seconds, resulting in a uniform ~0.3 mm PDMS thickness. The wafer was baked in a 70°C oven for at least 4 hours. Wells were cut using a custom razor blade stencil on a hard steel plate, and both sides were spray-washed with ethanol, then air-dried. The device had dimensions of 42 x 34 x 5 mm, and a 20 x 6.8 x 0.3 mm PDMS microwell array was attached as the floor of the main bath using liquid-tight silicon adhesive and a glass coverslip. Two channels for 0.5 mm diameter platinum wire electrodes (95% platinum, 5% ruthenium, Hoover & Strong, North Chesterfield, VA) ran the length of the bath and were used to field-stimulate cells at 0.5 or 1 Hz. A temperature probe and four 3 Ω power resistors along both sides of the bath were used to maintain an internal bath temperature of 36±1°C. Inflow and outflow channels with barbed fittings allowed for superfusion with Tyrode’s solution driven by external syringe pumps at 0.4 mL min^−1^. At this rate, the bath volume was exchanged every 2.3 min.

### Device chassis preparation

The chassis of the device was printed using a resin-cure lithographic printer (Form 2, Formlabs, Somerville, MA) and washed in an isopropanol bath for 20 min. Compressed air was used to clear all channels to prevent debris and resin build-up, and the device was allowed to dry for at least one hour. After the coverslip with PDMS microwells was secured onto the bottom of the device, the device was plasma treated with oxygen for 90 seconds in preparation for open-faced cell seeding.

### Functional testing of cardiomyocytes

Cardiomyocytes were imaged on a custom microscope apparatus in Tyrode’s solution (containing in mM: 140 NaCl, 5.4 KCl, 1.8 CaCl_2_, 1 MgCl_2_, 25 Hepes, and 10 glucose; pH 7.3). Prior to study, cells were incubated in darkness for 10 min with Tyrode’s solution supplemented with 2.2 μM Fura-2AM (EMD Millipore, Burlington, MA) and with Pluronic F-127 (0.022%, Sigma-Aldrich, St. Louis, MO, USA) for Ca^2+^ fluorescence imaging (except for dye-free cardiomyocytes in Fig. 3). After loading, cells were resuspended in fresh Tyrode’s solution and allowed to settle naturally until imaging. Cardiomyocyte Ca^2+^ transients and unloaded shortening contractions were measured using an inverted microscope (Eclipse Ti-U, Nikon, Tokyo, Japan) equipped with either the traditional or PDMS microwell perfusion bath. The baths were temperature-controlled (Cell MicroControls, Norfolk, VA) and continuously perfused with 36±1°C Tyrode’s solution. Cells were field-stimulated at 1 Hz for comparison between traditional and PDMS microwells, and at 0.5 Hz for all other experiments to reduce effects from experimental rundown. Cells were preconditioned under electrical stimulus and flow for >12 min before collecting functional records. Contractile events were imaged in real time by using a sarcomere-length camera system (HVSL, Aurora Scientific, Aurora, ON, Canada). Ca^2+^ transient measurements were recorded by using a low-pass filtered illumination with oscillating excitation wavelengths of 340 and 380 nm controlled by a RatioMaster fluorescence system (Horiba, Kyoto, Japan) with wavelengths switched every 10 ms. Fura-2 photobleaching over many repeat excitation cycles was compensated for using an empirically-derived linear correction. Data signals were recorded with a DAP5216a data acquisition system (Microstar Laboratories, Redmond, WA) and processed with custom software written in MATLAB (MathWorks, Natick, MA). Only rod-shaped cardiomyocytes with well-defined sarcomere striations were measured. Cardiomyocytes that did not respond to pacing were excluded regardless of appearance. From each recording, peak sarcomere length shortening, time to peak shortening (TTP), time from peak shortening to 50% re-lengthening (RT50), diastolic sarcomere length, magnitude of Ca^2+^ transient, and Ca^2+^ decay constant were calculated.

### Drug treatments

Cells in the drug comparison experiments were preconditioned for >12 min before perfusion was switched to either a drug-containing or control solution. The ‘Control’ group of cells was perfused with Tyrode’s solution only. The ‘DMSO’ group of cells was treated with Tyrode’s solution supplemented with 0.1% dimethyl sulfoxide (DMSO), the same concentration as the vehicle in the drug-containing solutions. The ‘Dobutamine’ group of cells was treated with Tyrode’s solution supplemented with 40 nM of dobutamine (hydrochloride) (Cayman Chemical, Ann Arbor, MI) carried in 0.1% DMSO. The ‘Omecamtiv Mecarbil’ group of cells was treated with Tyrode’s solution supplemented with 150 nM CK-1827452 (Cayman Chemical, Ann Arbor, MI) carried in 0.1% DMSO. Each group of cells was constantly superfused with each respective solution after preconditioning until the completion of the experiment.

### Single-cell RT-qPCR

After completing functional testing, individual cardiomyocytes were immediately aspirated and deposited into separate microcentrifuge tubes containing SingleShot Cell Lysis buffer (Bio-Rad, Hercules, CA) supplemented with DNase, and proteinase K. Aliquots were frozen at −80°C overnight to aid in cell lysis, then heated to 75°C for 5 min to digest gDNA. cDNA was synthesized using the iScript Advanced cDNA Synthesis Kit (Bio-Rad, Hercules, CA), then preamplified using SsoAdvanced PreAmp Supermix (Bio-Rad, Hercules, CA) to amplify target cDNA yield. A nested primer design was implemented to reduce nonspecific binding. qPCR was performed using SsoAdvanced Universal SYBR Green Supermix on a CFX Connect Detection System (Bio-Rad, Hercules, CA). Genes of interest were GAPDH, PRKACA, PPP1CA, PPP2CB, and ADRB2. ADRB1 was excluded because a nested primer design could not be generated. Transcript abundance was normalized to GAPDH, then to the respective transcript adbundance in control cells. All primer sequences are listed in the Supplemental Material.

### Statistical analysis

Data were analyzed using Prism (GraphPad Software, San Diego, CA), Minitab (Minitab Inc., State College, PA), and MATLAB (MathWorks, Natick, MA). All data were presented as mean ± 95% confidence interval. Comparisons were conducted via one-way ANOVA test followed by multiple comparisons (Kurskal-Wallis post hoc analysis), or via an unpaired two-tailed Student’s t-test. Significance was defined by p < 0.05. For comparison of variance, Levene’s test (assuming non-Gaussian distributions) was used.

## RESULTS

To assess whether the PDMS microwell array had any significant effect on the integrity of the cardiomyocytes, sarcomere shortening dynamics, or Ca^2+^ transient measurements, we compared the PDMS microwell device (n = 57 cells) to a more traditional setup in which cardiomyocytes are deposited directly onto a coverglass that forms the bottom of the superfusion bath (n = 58 cells). After allowing cell aliquots to precondition for at least 12 min under 1 Hz pacing, one set of sarcomere length shortening and Ca^2+^ transient records were collected from each cell within the time frame of 45 min after the start of pacing. Bright-field micrographs taken before and after 5 min of perfusion highlight cell drift in the traditional setup, which is one of the key limitations for capturing multiple functional records of cardiomyocytes over time using a traditional setup (Fig. 2A). By contrast, cell loss or drift was not observed in the microwell device.

**Figure 2:**
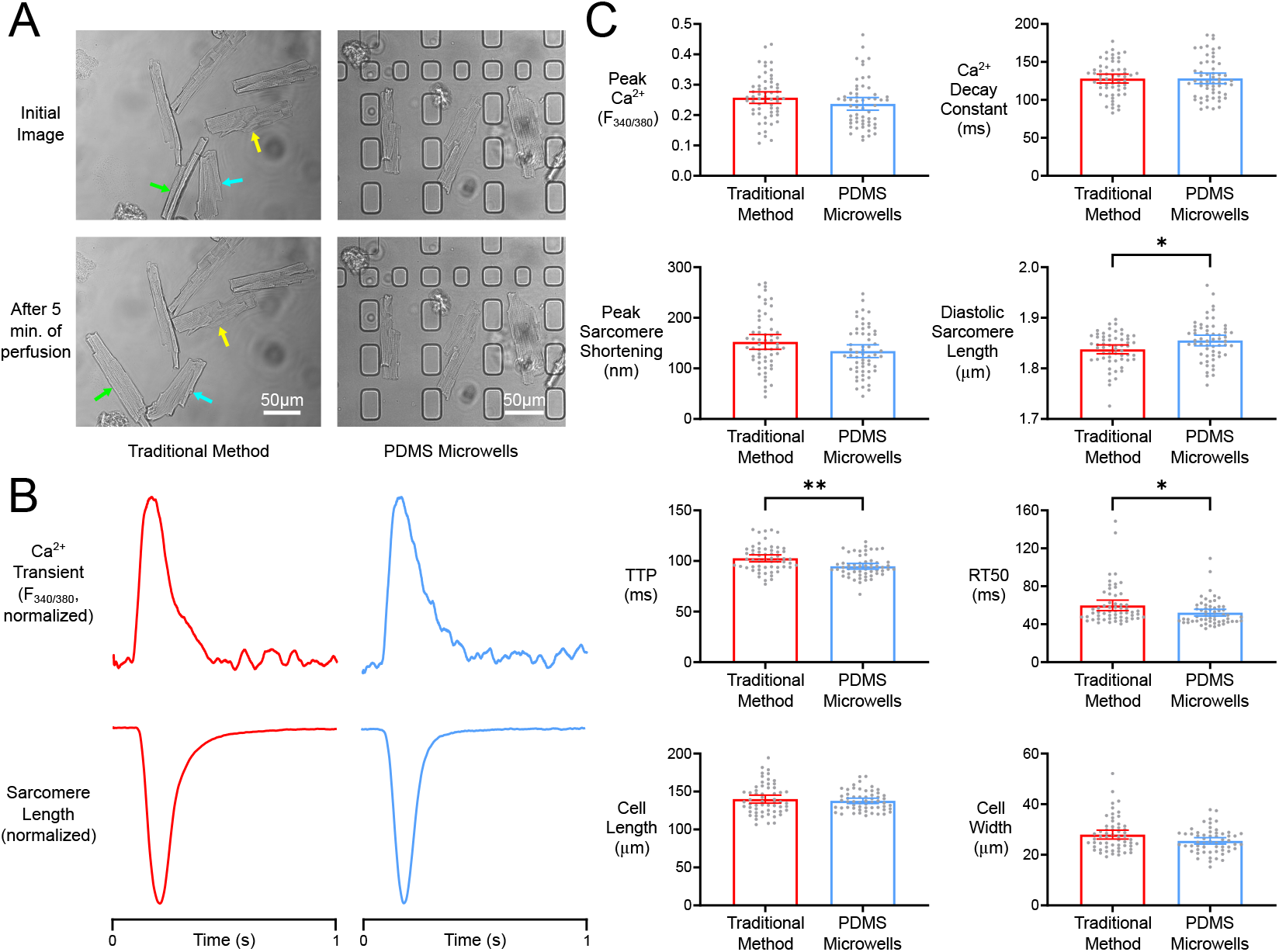
Comparison of traditional and PDMS microwell methods. *(A)* Bright-field micrographs of cardiomyocytes at 30x magnification in traditional bath (left) and PDMS microwells (right) before and after a 5 min interval to demonstrate cell drifting. Colored arrows indicate the same cells at different time points. *(B)* Representative normalized Ca^2+^ transients and sarcomere shortening using traditional bath (left) and PDMS microwells (right). *(C)* Comparison of cell properties between traditional and PDMS microwell methods. Statistical significance of means indicated by * (p < 0.05) and ** (p < 0.001).

Representative Ca^2+^ transients and sarcomere shortening records show the overall similarity between measurements made on the respective devices (Fig. 2B). No significant differences between the mean Ca^2+^ release magnitude, Ca^2+^ decay constant, peak sarcomere shortening, cell length, or cell width were detected (Fig. 2C). Both time to peak shortening (TTP, p < 0.001) and time from peak shortening to 50% re-lengthening (RT50, p < 0.05) were reduced in the cells measured in PDMS microwells. Resting sarcomere length was shown to be slightly longer in cells tested in the new device (p < 0.05). Although statistically significant, the observed mean differences between the two methods were relatively small (~9%). If anything, the larger mean resting sarcomere length of cells in the PDMS microwells may indicate beneficial effects of shielding cells from direct fluid shear, resulting in slightly better cardiomyocyte viability.

We next sought to establish the baseline variability of cardiomyocyte contractile behavior during experiments lasting many tens of minutes. This inherent variability over time is the backdrop against which drug responses occur. The behavior of six cells was followed over the course of one hour with repeated measurements made roughly every two minutes. To investigate the influence of the Fura-2 Ca^2+^ indicator on long-term cell dynamics, we tracked three cells loaded with indicator and three cells without. The cells were stimulated at 0.5 Hz to limit the effect of repeated pacing events over a long period of superfusion. Ca^2+^ transients and sarcomere shortening records of a Fura-2-loaded cell over five time-points after preconditioning show no obvious rundown or degradation of cell integrity (Fig. 3A).

**Figure 3:**
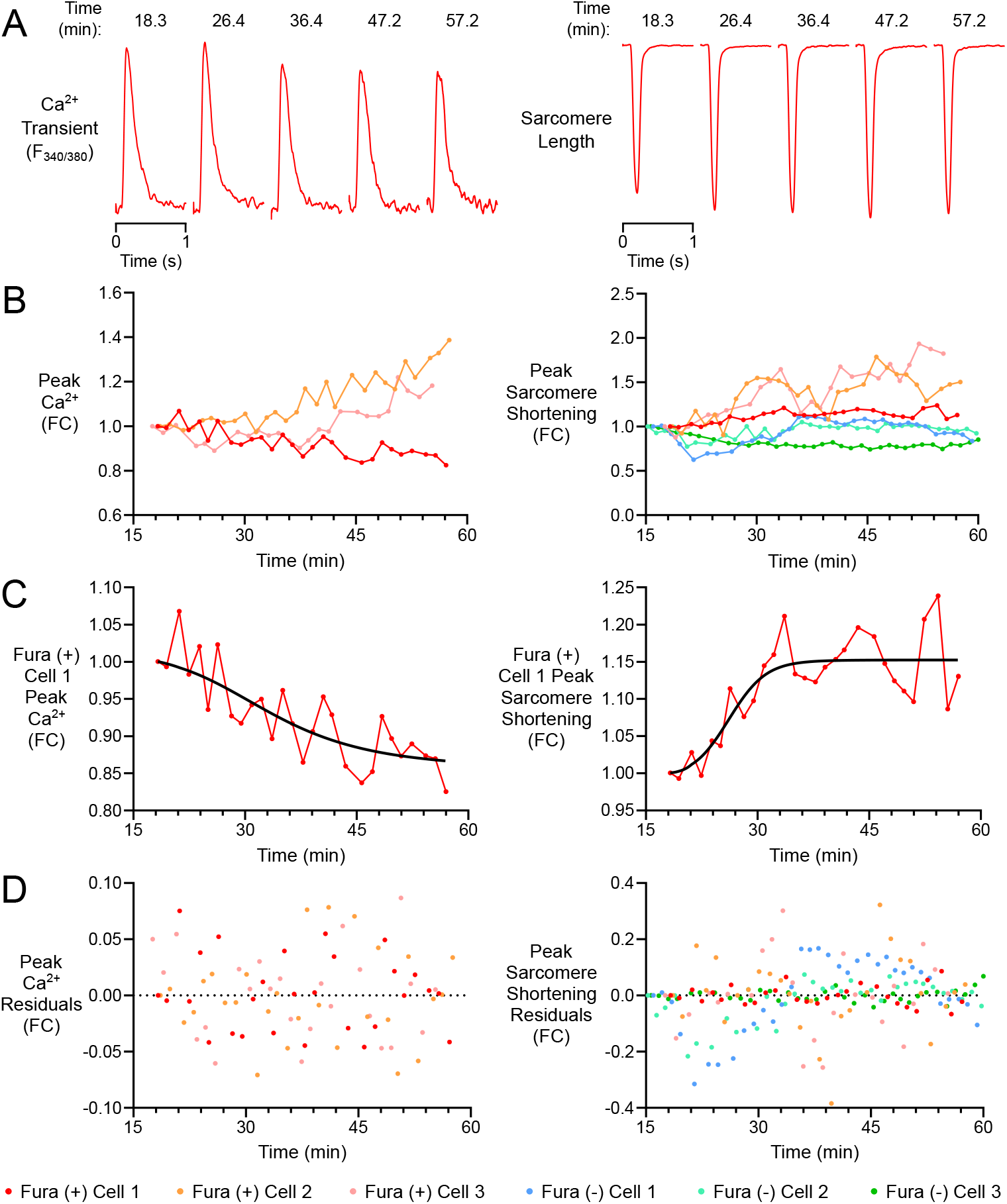
Variability of single cells with and without Fura-2 measured over one hour. *(A)* Ca^2+^ transients and sarcomere lengths of a Fura-loaded cell (Fura(+) Cell 1) at five time-points over one hour. *(B)* Normalized peak Ca^2+^ and peak sarcomere shortening of Fura-loaded and non-Fura-loaded cardiomyocytes over one hour, expressed as a fold change (FC) compared to the initial record after preconditioning. *(C)* Hill-fit regressions of Fura(+) Cell 1 peak Ca^2+^ and peak sarcomere shortening. *(D)* Residual values from hill-fit regressions of each cell.

Properties such as peak Ca^2+^ release and peak sarcomere shortening were calculated from measurements at each time-point. Comparing these values to those that manifested in the same cell at the initiation of the experiment allows the fold change (FC) of the various functional metrics to be represented over time (Fig. 3B). Each cell’s FC values over time for a given parameter were fit with a Hill regression to summarize time-based trends in the data as the cells reached a dynamic steady state (Fig. 3C). The deviation of FC values from the Hill equation (the residuals) of each cell indicate that both Fura-2-loaded and dye-free cells fluctuate continuously over the entire course of the experiment (Fig. 3D). These fits had an average root mean square error (RMSE) of 0.037 FC for peak Ca^2+^ transient amplitude and 0.102 FC for peak sarcomere shortening in the Fura-2-loaded cells, with an average RMSE of 0.075 FC for peak sarcomere shortening in dye-free cells. Fura-2 loading did not appear to affect the variation of peak sarcomere shortening, diastolic sarcomere length, TTP, or RT50 over time based on a comparison of the RMSE means found in the Supplemental Material. Full records of TTP, RT50, diastolic sarcomere length, and Ca^2+^ decay constant fold changes are included in the Supplemental Material.

Having established viability of cells in the microwell device and characterized their stability during prolonged experiments, we next used the device to quantify the responses of several individual cells to dobutamine and omecamtiv mecarbil (OM). Cardiomyocytes were superfused with either drug (40 nM dobutamine, n = 41 cells; 150 nM OM, n = 34 cells) for up to one hour after 12 min of preconditioning. For control groups, cells were also superfused with Tyrode’s solution alone (Control, n = 19 cells), as well as a vehicle-only control group superfused with 0.1% dimethyl sulphoxide in Tyrode’s (DMSO, n = 17 cells). Representative Ca^2+^ transients and sarcomere shortening records illustrate the mean behavior of cardiomyocytes under the various test conditions (Fig. 4A). Gray insets within the sarcomere shortening records show the unadjusted transients to emphasize differences in diastolic sarcomere length. Eight representative time-course records of the changes in Ca^2+^ and contractile properties during drug treatment are shown in Fig. 4B. These responses show the positive inotropic effects of both dobutamine and OM. They also suggest substantial heterogeneity in the responses of individual cells.

**Figure 4:**
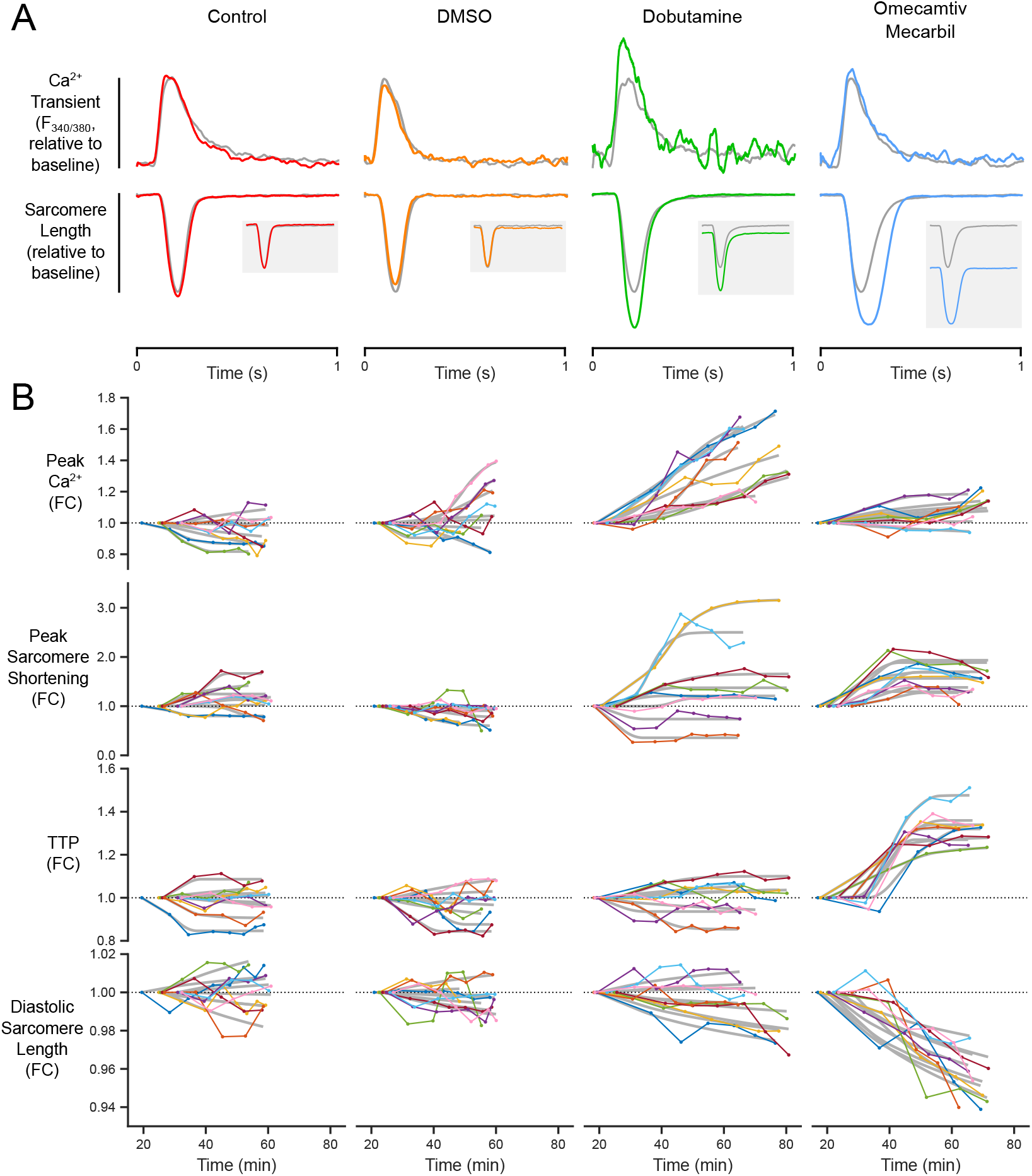
Effects of dobutamine and omecamtiv mecarbil on functional properties. *(A)* Representative Ca^2+^ transients and sarcomere shortening records from each treatment group after preconditioning (gray) and after drug exposure (colored). Both are scaled to maintain the same peak sarcomere shortening values between the pre-treatment transients. Sarcomere shortening records are aligned at the same diastolic sarcomere length. Gray subplots show both shortening records unaligned. *(B)* Eight representative time-courses of functional parameters for control (n = 19), 0.1% DMSO control (n = 17), dobutamine (n = 41), and OM (n = 34) groups. Hill fit regressions are shown in gray behind the time-course data. Columns are labeled as in panel A.

As before, the time-courses of cellular properties were fitted with a Hill regression in order to describe their dynamic steady-state responses (Fig. 4B, gray lines). FC residuals from Hill fits were similar for the most part across all treatment groups, indicating that time variation in cellular behavior was not affected by the drugs studied (comparisons of RMSE between groups can be found in the Supplemental Material). From these fits, steady-state FC values for Ca^2+^ and contractile properties were aggregated for each treatment and compared via a one-way ANOVA (Fig. 5). On average, dobutamine caused increases in Ca^2+^ release (p < 0.0001) and peak shortening (p < 0.05) with little change in kinetics. OM also increased the magnitude of sarcomere shortening (p < 0.01), but with a concomitant lengthening of TTP (p < 0.0001) and RT50 (p < 0.0001), and a large decrease in the diastolic sarcomere length (p < 0.0001). The variance of steady-state response properties was also assessed and compared between dobutamine- and OM-treated groups. Peak Ca^2+^ release (p < 0.0001), Ca^2+^ decay constant (p < 0.05), and peak sarcomere shortening (p < 0.05) responses were all significantly more variable in dobutamine-treated cardiomyocytes compared to OM (Levene’s Test).

**Figure 5:**
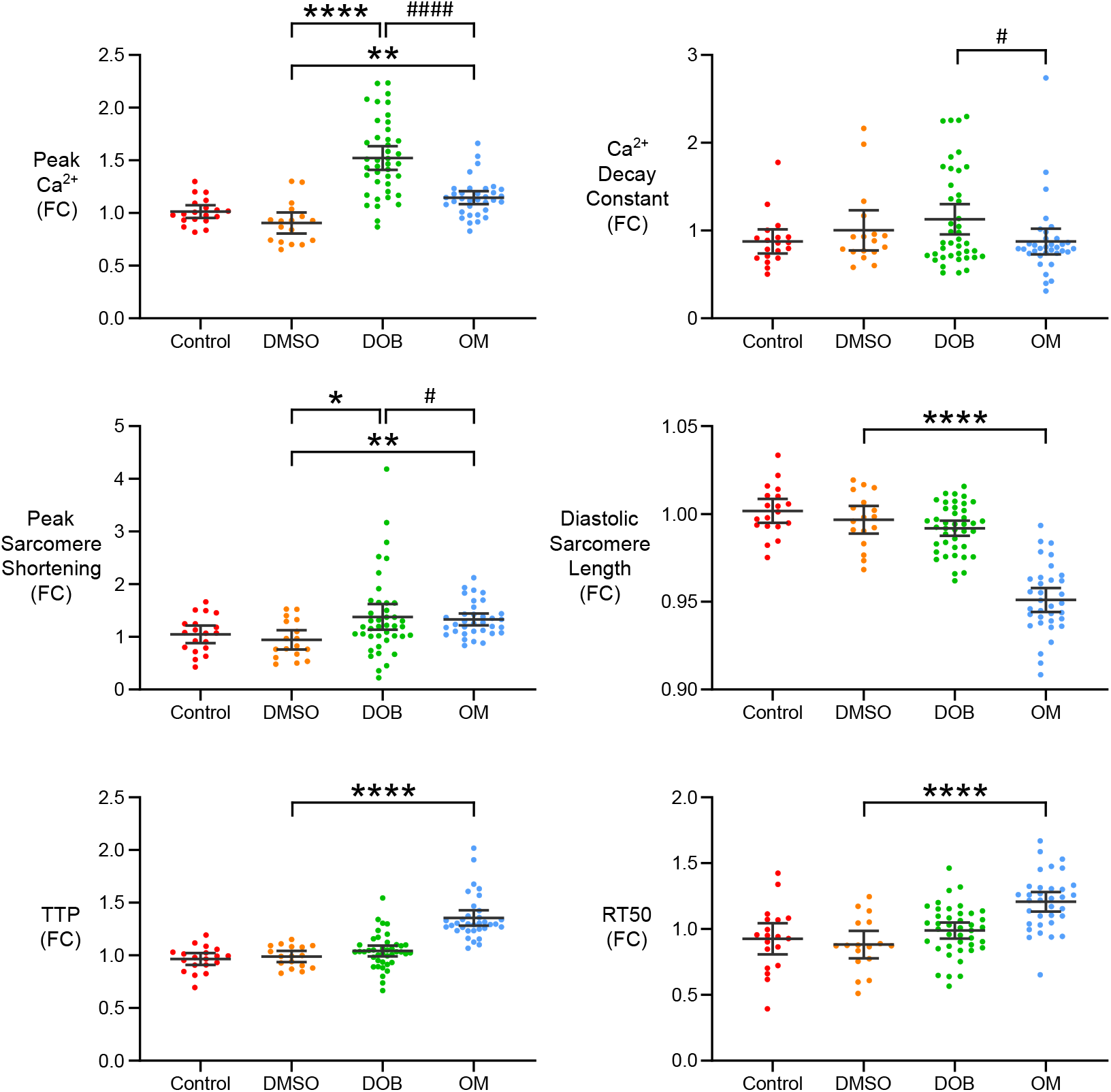
Summary of functional parameter changes after drug treatment. Functional parameter values determined from the maximum of hill fit regression. Statistical significance between means represented by * (p < 0.05), ** (p < 0.01), **** (p < 0.0001). Statistical significance between variances represented by # (p < 0.05), #### (p < 0.0001). ‘DOB’ is an abbreviation of dobutamine.

Dobutamine-treated cardiomyocytes also exhibited a surprising dissociation between increases in peak Ca^2+^ release and increases in peak sarcomere shortening. This is clearly seen in the records of two dobutamine-treated cells (Fig. 6A). In the first case, the drug causes a robust increase in Ca^2+^ release while sarcomere shortening actually decreases. In another cell, the Ca^2+^ transient is unchanged even as sarcomere shortening doubles. These stark contrasts in response are still clearer when visualized as changes in each cell’s Ca^2+^-sarcomere shortening hysteresis loop (Fig. 6C). Examining FC in steady-state Ca^2+^ release and peak sarcomere shortening for each of the dobutamine-treated cells shows that some form of mismatch is present in the majority of cells (Fig. 6C). Furthermore, 16 of 41 cells showed either no change or a decrease in contractility after dobutamine treatment.

**Figure 6:**
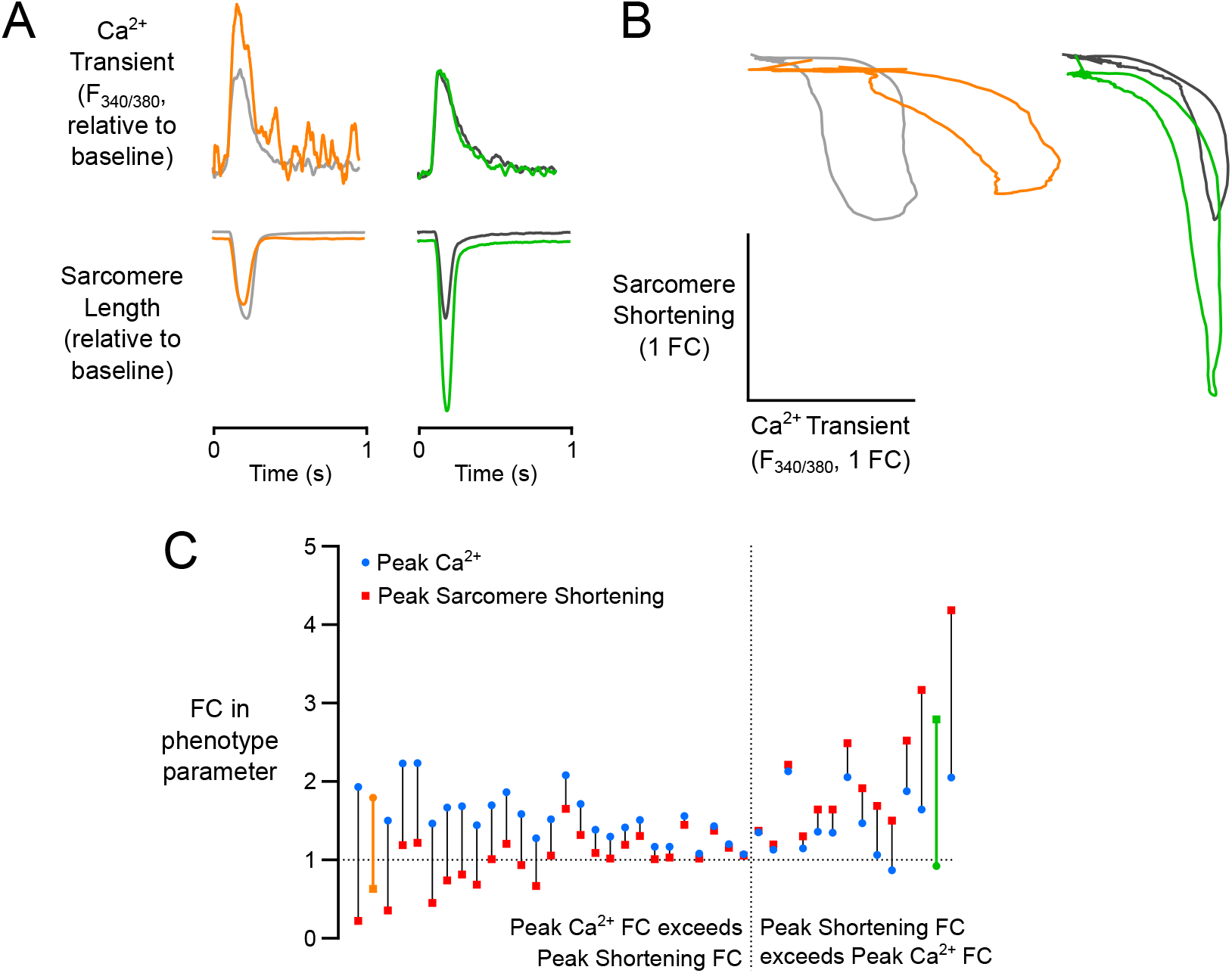
Relative changes in peak Ca^2+^ and peak sarcomere shortening in dobutamine-treated cardiomyocytes. *(A)* Representative Ca^2+^ transients and sarcomere shortening records of a cardiomyocyte that displayed a positive change in Ca^2+^ release with a negative change in peak sarcomere shortening (left) and a cell that displayed no change in Ca^2+^ release, yet a positive change in peak sarcomere shortening (right). Gray transients indicate records taken immediatley after preconditioning, while colored transients were taken after sustained dobutamine treatment. Both are scaled to maintain the same peak sarcomere shortening values between the pre-treatment transients. *(B)* Ca^2+^-sarcomere shortening hysteresis loops of each of the select transients from panel A emphasize the uniaxial effects of dobutamine on each cell. *(C)* Each data pair corresponds to a single cell’s fold change response for peak Ca^2+^ and peak sarcomere shortening, rank-ordered by the difference in fold change between the two functional properties. Cells from panels A & B are highlighted with their respective colors.

This finding reinforced our original hypothesis concerning the variability of cardiomyocyte responses to *β*-adrenergic stimulus. It also underscored the likelihood that the abundance of gene transcripts associated with this pathway would differ significantly among cardiomyocytes. Accordingly, we resolved to examine the transcriptomic activity of individual cells and attempt to correlate this information with their particular responses to dobutamine.

A new batch of cardiomyocytes was superfused with 40 nM dobutamine for 35 min while monitoring cellular responses (n = 7). The magnitudes of Ca^2+^ and shortening responses were used to rank-order cells as before (Fig. 7A). After functional measurements were taken, each cell was individually aspirated and deposited into its own microcentrifuge tube containing cell lysis buffer. Cell lysates underwent reverse transcriptase followed by a preamplification step to amplify our genes of interest: ADRB2 (*β*2 adrenergic receptor), PRKACA (protein kinase A, PKA), PPP1CB (protein phosphatase 1), PPP2CA (protein phosphatase 2), and GAPDH (housekeeping). PKA phosphorylates regulatory Ca^2+^ and sarcomeric proteins as a result of *β*-adrenergic stimulation. Protein phosphatase 1 & 2 dephosphorylate those same proteins. Real time PCR was used to quantify abundance of each transcript relative to GAPDH, then compared to the transcript abundance in two untreated control cells. ADRB2 was shown to be downregulated across all dobutamine-treated cells compared with untreated control cells (p < 0.01; Fig. 7B). This is a known result of sustained *β*-adrenergic stimulation and demonstrates that gene expression 35 min after dobutamine treatment already reflects an adaptive cellular response. We subsequently probed the coupled physiological and gene expression responses of individual cells to dobutamine in search of properties that were significantly correlated. This analysis yielded a significant negative correlation between PRKACA expression and Ca^2+^ release FC following dobutamine application (p < 0.05; Fig. 7C). In other words, the data suggest upregulation of PKA in cells that had a decrease in Ca^2+^ release following dobutamine, and a downregulation of PKA in cells that experienced increased Ca^2+^ release.

**Figure 7:**
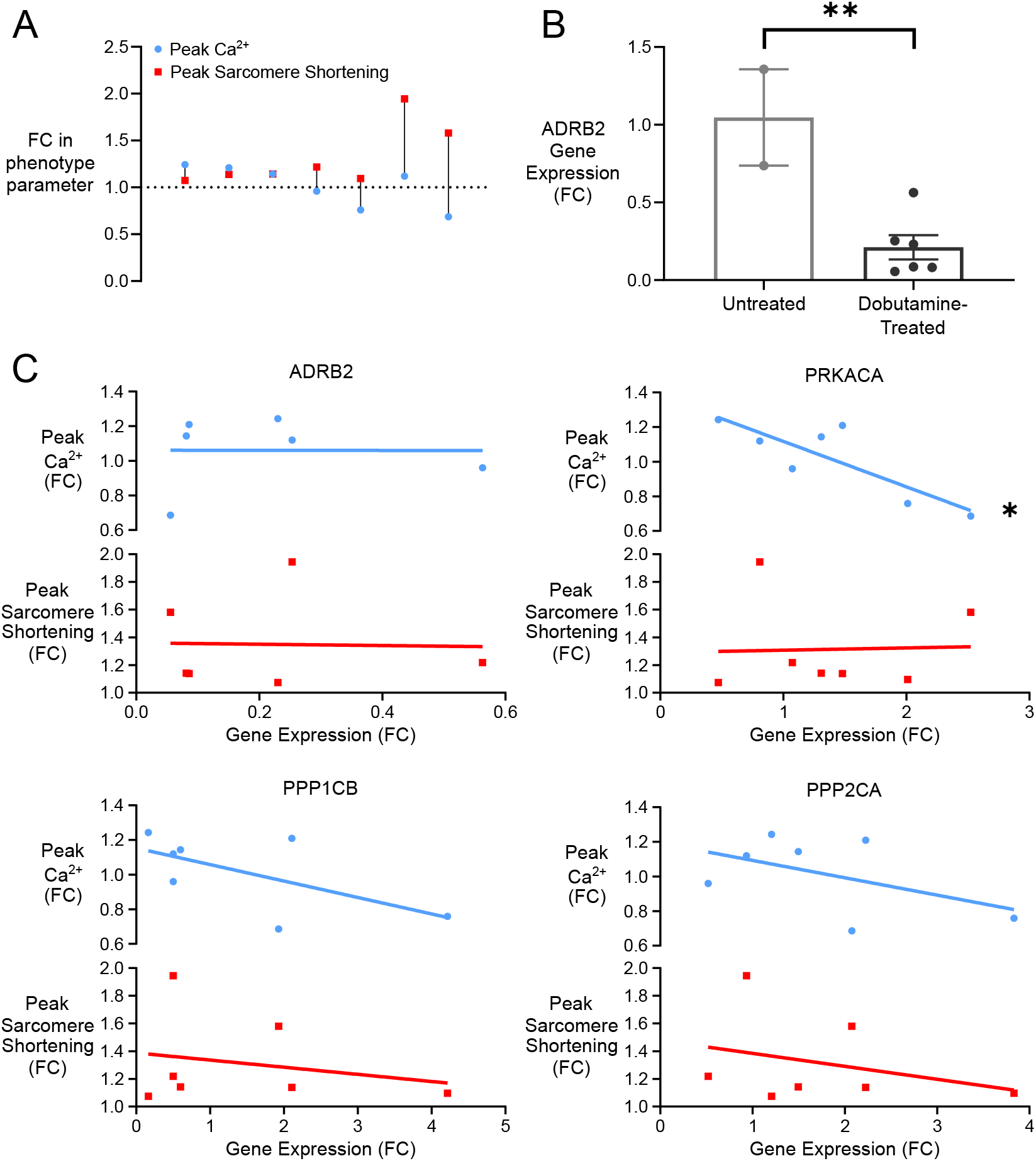
Transcriptomic analysis of dobutamine-treated cells correlated with fold change in functional properties. *(A)* Each data pair corresponds to a single cell’s fold change response for peak Ca^2+^ and peak sarcomere shortening, rank-ordered by the difference in fold change between the two functional properties. *(B)* Comparison of expression of *β*2-adrenoceptor in control and dobutamine-treated cells. Significance in mean represented by ** (p < 0.01). *(C)* Gene expression fold changes correlated with both peak Ca^2+^ and peak sarcomere shortening fold changes. Significant correlation represented by * (p < 0.05).

## DISCUSSION

This study comes at a time of increased interest in characterizing the functional behavior and gene expression of individual cardiomyocytes. For instance, micro-patterned surfaces have been employed to generate large arrays of single induced pluripotent stem cell (iPSC) derived cardiomyocytes (4, 5). These arrays allow a variety of functional measurements to be performed on many individual cells. Others have succeeded in relating functional measurements of individual iPSC-derived cardiomyocytes with cell-specific myosin heavy chain isoform expression (6). The microwell capture device described here is a technical advance that provides new and useful ways to link isolated adult cardiomyocyte function with gene expression. We have also demonstrated the ability of the device to quantify cell-to-cell variation in sustained drug responses.

Our initial objective was to test the hypothesis that an inotrope operating via multiple intracellular intermediaries (dobutamine) would produce more variable responses than one targeted to a specific molecule (OM). This was clearly supported by the data (Fig. 5) and suggests the presence of substantial noise in the expression of genes that participate in the *β*-adrenergic signaling pathway. Cardiomyocyte responses to OM were consistent with other recent reports (3, 7), including the observation that OM increases contractility at the expense of diastolic function.

Perhaps the most striking finding in this study is the frequent occurrence of discordant Ca^2+^-contraction coupling responses in cardiomyocytes following dobutamine administration (Fig. 6). Because intracellular Ca^2+^ is the primary signal regulating cardiomyocyte contraction, these two properties are expected to be tightly correlated. However, we observed several examples of dobutamine-treated cardiomyocytes that experienced decreased contractility in spite of elevated Ca^2+^ release, and others that showed large increases in contractility with no corresponding increase in Ca^2+^.

These seemingly paradoxical responses are less surprising when the complexity of the *β*-adrenergic signaling pathway and the diversity of its subcellular targets are considered. At the level of the intact myocardium, *β*-adrenergic stimulus causes both increased contractility (inotropy) and enhanced relaxation rate (lusitropy) (8). This highly desirable combination of increased strength and speed is accomplished through a multitude of simultaneous subcellular changes. On one hand, Ca^2+^ release is augmented via PKA phosphorylation of L-type Ca^2+^ channels and other targets (9). PKA also phosphorylates myosin binding protein C (MyBP-C) which primarily increases the rate of crossbridge binding (10). All of these events tend toward increases in the magnitude and duration of contraction. On the other hand, PKA phosphorylates phospholamban and troponin I (TnI). These events promote relaxation and tend to reduce force production (11–13). When enacted together, these apparently antagonistic effects combine to produce simultaneous increases in inotropy and lusitropy.

The presence of antagonistic subcellular processes that are all activated by the *β*-adrenergic signaling cascade raises the possibility of extreme cell behavior if the opposing effects are not perfectly balanced. We believe that this explains the diversity of dobutamine responses revealed in our experiments (Fig. 6). For instance, if a cell were randomly deficient in regulatory proteins that anchored PKA specifically to MyBP-C, such as myomegalin (14), it might display the expected increase in Ca^2+^ release but fail to show increased contractility. PKA could still be localized to the myofilament by other anchoring proteins such as troponin T (15), but the Ca^2+^-desensitizing effects of TnI phosphorylation (12) would operate with less opposition. Interestingly, such scenarios were contemplated by Negroni et al., who constructed an integrative cardiomyocyte model representing the various effects of *β*-adrenergic signaling on electrical, Ca^2+^ handing, and contractile systems of the cell (13). In order to illustrate the importance of each phosphorylation target on achieving combined inotropic and lusitropic effects, they simulated cardiomyocyte Ca^2+^ transients and unloaded shortening while eliminating each target in turn. Their simulations show dramatic shifts away from the expected behavior when one or more components of the response are suppressed. In fact, the diversity of functional outputs they were able to predict is strikingly similar to the modes of discordant Ca^2+^-contraction responses we observed among measured single-cell responses.

In spite of a high degree of individual variation, on average the cardiomyocytes exposed to dobutamine behaved in the expected manner. Therefore, it is likely that when coupled electrically and mechanically in the intact myocardium, *β*-adrenergic response heterogeneity is of minimal consequence. Nevertheless, the existence of this heterogeneity is meaningful in at least two ways. First, it provides evidentiary support for the premise advanced by Negroni et al., namely that each of the disparate sub-cellular functions targeted by *β*-adrenergic signaling is critical to its physiological role of simultaneously enhancing ventricular inotropy and lusitropy. Second, when coupled with single-cell biochemical assays, cell-to-cell differences can be leveraged to discover new features of the *β*-adrenergic pathway that could be exploited for cardiac drug development.

In experiments that link aspects of the dobutamine response to gene expression, we established the technical feasibility of this approach and provide an example of how it can lead to unique insights. Single-cell gene expression profiling revealed that all dobutamine-treated cells responded by downregulating *β*2 receptor transcript within 35 minutes, conforming to expectations set by studies in other cell types (16, 17). Interestingly, the degree of receptor downregulation was not correlated with the strength of the functional response to dobutamine (Fig. 7C). The expression of genes encoding protein phosphatases that reverse PKA activity did not correlate with functional dobutamine response either. However, transcript abundance for the catalytic subunit of PKA (PRKACA) was negatively correlated with the dobutamine-stimulated increase in Ca^2+^ release.

One possible explanation of this finding would be a hypothetical feedback mechanism that monitors both receptor activity and expected functional changes (such as increased Ca^2+^ release). When receptor activity is detected in the absence of increased Ca^2+^ release, this pathway could rectify the mismatch by increasing expression of PKA. If receptor expression is tightly controlled by cells, it follows that intermediate *β*-adrenergic pathway components would also be subject to homeostatic control. Further insights will be possible as this approach is coupled to unbiased screens such as RNAseq. This process could uncover entirely new genes that modulate *β*-adrenergic signaling in cardiomyocytes.

It must be acknowledged that phenomena identified in isolated rat cardiomyocytes are not guaranteed to apply unambiguously to physiological situations. The process of isolation disconnects cardiomyocytes from the extracellular matrix and from neighboring cells, with which they share mechanical and electrical connections (18). Undoubtedly this perturbs cellular properties and may interfere with signaling microdomains critical to the *β*-adrenergic response (19, 20). For these reasons, it will ultimately be necessary to confirm findings obtained with this system in multi-cellular preparations.

## AUTHOR CONTRIBUTIONS

Microwell device conceived by JAC, JDW, and SGC. Experiments designed by JAC and SGC. Data collected by JAC and JDW. Data analyzed and manuscript written by JAC, JDW, and SCG.

## ACKNOWLEDGMENTS

The authors thank Adriel S. Sumathipala and Burak Dura for their contributions toward early prototypes of the PDMS microwells. This work was supported by a National Science Foundation Grant (CMMI-1562587) to S.G.C.

## SUPPLEMENTARY MATERIAL

**Figure 8:**
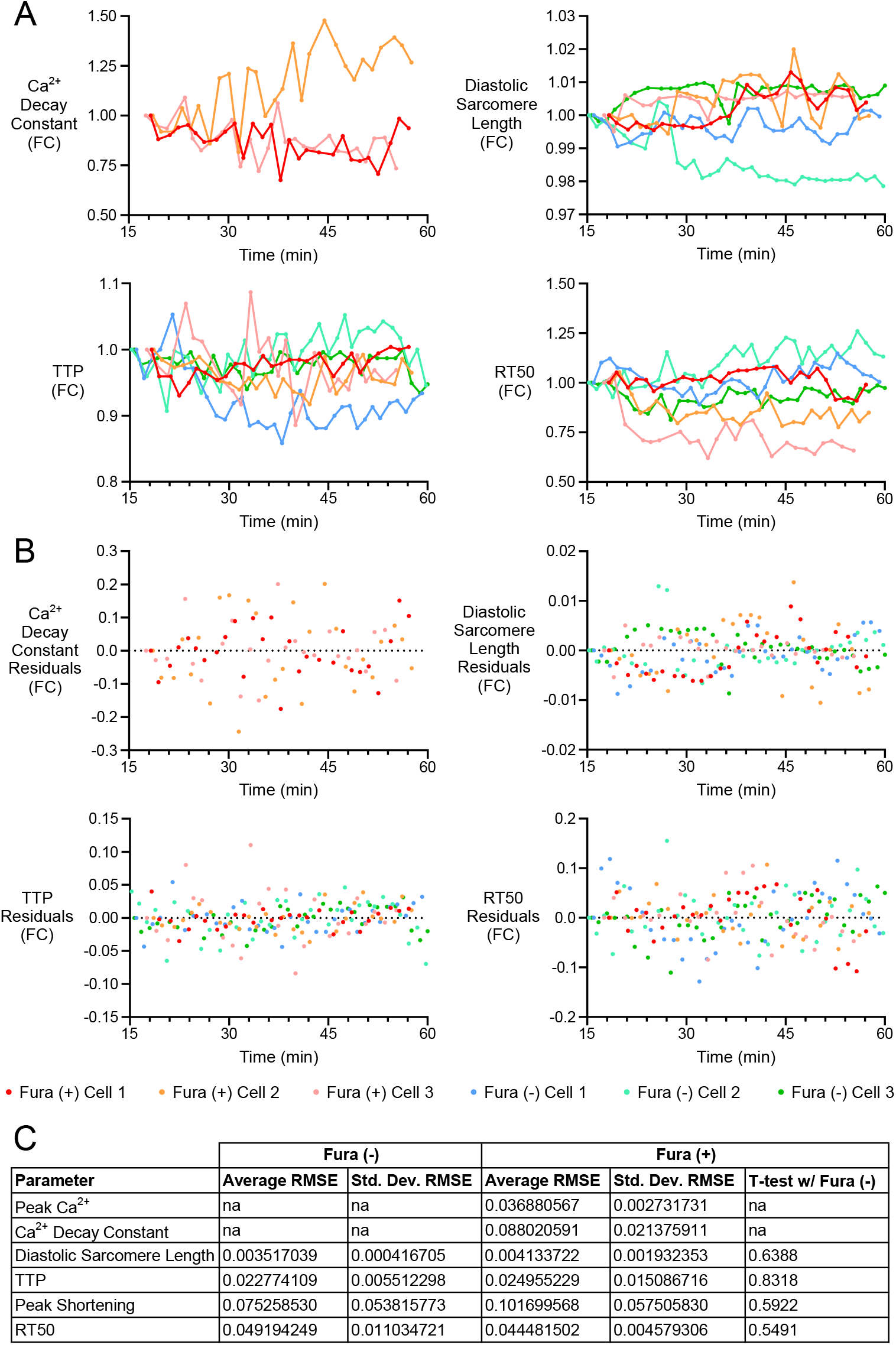
Single-cell time-course records and residuals. *(A)* Functional parameters of Fura-loaded and non-Fura-loaded cardiomyocytes over one hour *(B)* Residual values from hill-fit regressions of each cell. *(C)* Root mean square error (RMSE) comparisons between groups for each parameter.

**Figure 9:**
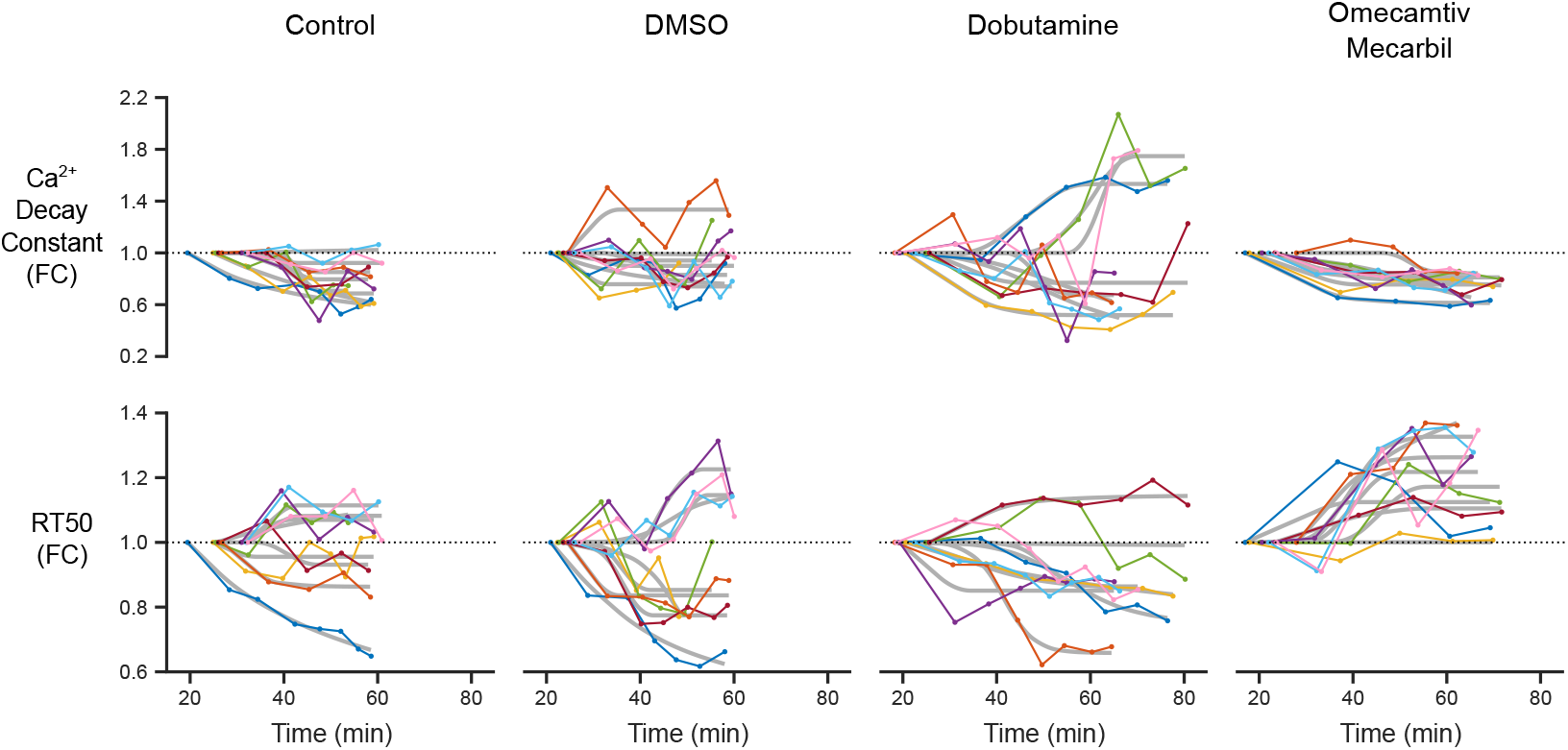
Ca^2+^ decay constant and RT50 of drug traces over time. Eight representative time-courses of functional parameters for control (n = 19), 0.1% DMSO control (n = 17), dobutamine (n = 41), and OM (n = 34) groups. Hill fit regressions are shown in gray behind the time-course data.

**Figure 10:**
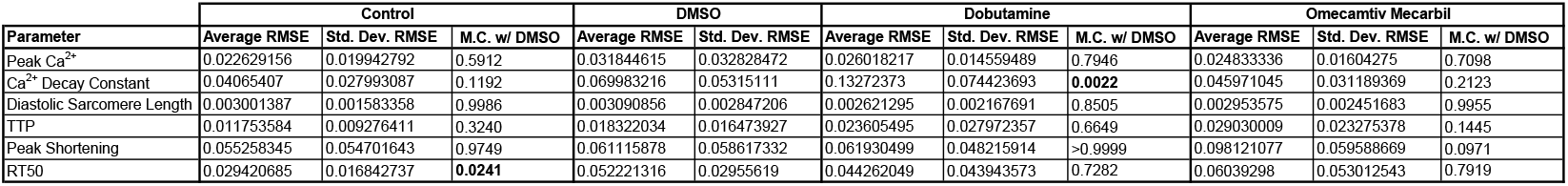
Root mean square error comparisons between drug hill fit regressions. Statistical significance between mean RMSE values from a post hoc multiple comparison to DMSO signified by bold p-values.

**Figure 11:**
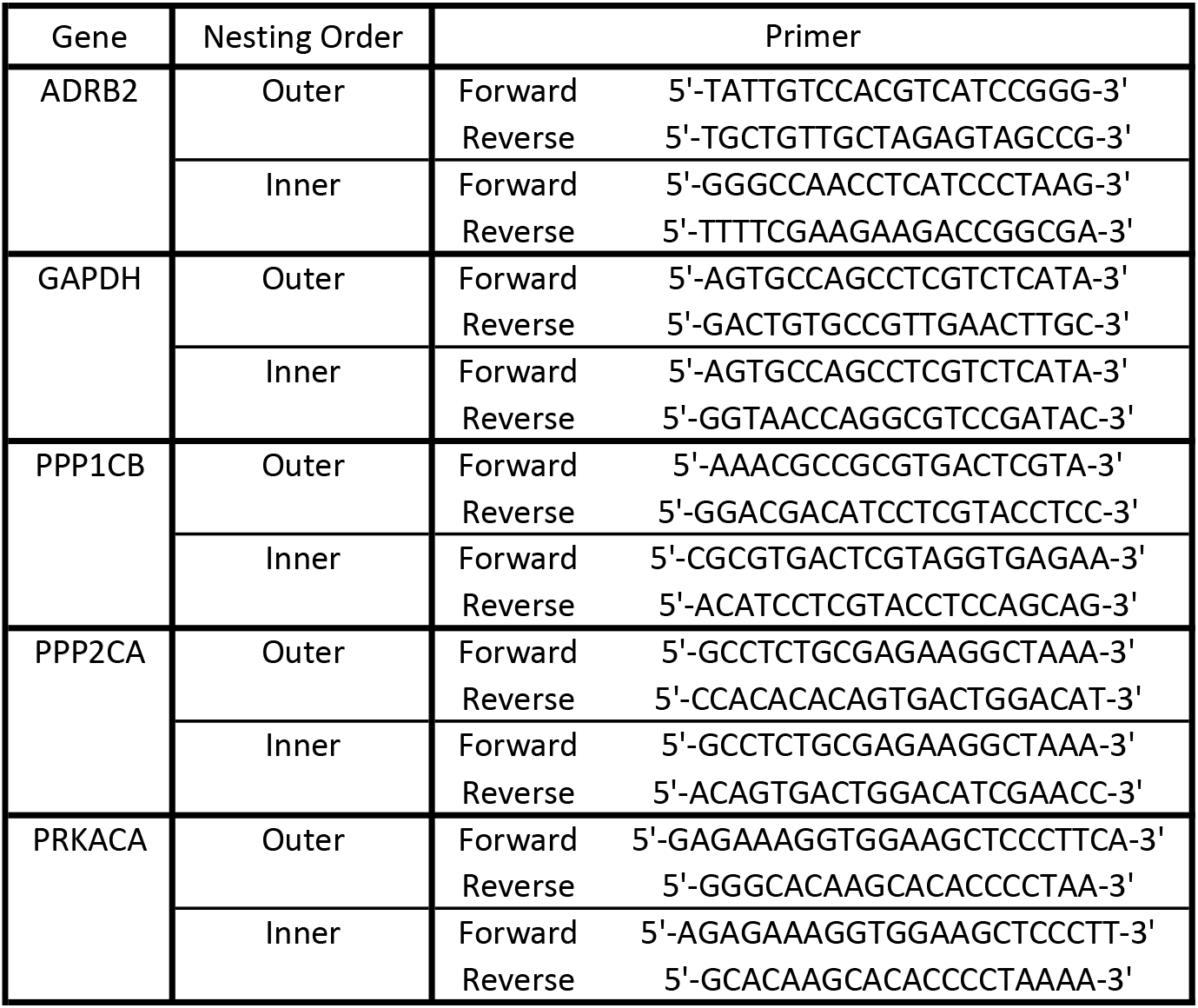
RT-qPCR Primers.

